# Tissue protective role of Ganetespib in SARS-CoV-2-infected Syrian golden hamsters

**DOI:** 10.1101/2022.12.27.521979

**Authors:** Luiz Gustavo Teixeira Alves, Morris Baumgardt, Judith Hoppe, Theresa C. Firsching, Julia M. Adler, Guido Mastrobuoni, Jenny Grobe, Katja Hönzke, Stefan Kempa, Achim D. Gruber, Andreas C. Hocke, Jakob Trimpert, Emanuel Wyler, Markus Landthaler

## Abstract

The emergence of new SARS-CoV-2 variants, capable of escaping the humoral immunity acquired by the available vaccines, together with waning immunity and vaccine hesitancy, challenges the efficacy of the vaccination strategy in fighting COVID-19. Improved therapeutic strategies are therefore urgently needed to better intervene particularly in severe cases of the disease. They should aim at controlling the hyper-inflammatory state generated upon infection, at reducing lung tissue pathology and endothelial damages, along with viral replication. Previous research has pointed a possible role for the chaperone HSP90 in SARS-CoV-2 replication and COVID-19 pathogenesis. Pharmacological intervention through HSP90 inhibitors was shown to be beneficial in the treatment of inflammatory diseases, infections and reducing replication of diverse viruses. In this study, we analyzed the effects of the potent HSP90 inhibitor Ganetespib *in vitro* on alveolar epithelial cells and alveolar macrophages to characterize its effects on cell activation and viral replication. Additionally, to evaluate its efficacy in controlling systemic inflammation and the viral burden after infection *in vivo*, a Syrian hamster model was used. *In vitro*, Ganetespib reduced viral replication on AECs in a dose-dependent manner and lowered significantly the expression of pro-inflammatory genes, in both AECs and alveolar macrophages. *In vivo*, administration of Ganetespib led to an overall improvement of the clinical condition of infected animals, with decreased systemic inflammation, reduced edema formation and lung tissue pathology. Altogether, we show that Ganetespib could be a potential medicine to treat moderate and severe cases of COVID-19.

## Introduction

The coronavirus disease 2019 (COVID-19) is a global threat with more than 600 million people infected and more than 6 million deaths reported worldwide since the beginning of the pandemic. Estimations by the WHO, based on the excess of average death rates, consider even higher numbers, counting up to 15 million deaths^1,2^. Infections with the beta-coronavirus SARS-CoV-2, the causal agent of COVID-19, lead mostly to rather mild symptoms or remain asymptomatic. However, in around 5% of the cases, the disease can progress to severe acute respiratory distress syndrome (ARDS), with increased lung injury, pulmonary barrier dysfunctions and edema formation in the lungs, being associated with high mortality rates^3,4^. The mechanisms involved in its pathogenicity are still incompletely understood. Dysregulated systemic and mucosal immune responses, elevated pro-inflammatory cytokine levels, increased neutrophil recruitment, and robust activation of platelets and the coagulation cascade, are frequently observed manifestations of the disease^5–14^. Pre-existing comorbidities, such as diabetes, obesity and cardiovascular disease or age were correlated to higher chances of developing more severe cases of the disease and higher mortality rates. This is probably due to an impaired immunological response to the virus^10,15^. SARS-CoV-2 cellular entry occurs after binding of the viral spike protein to the angiotensin converting enzyme 2 (ACE2), expressed on the surface of different cell types. Within the lower airways, pulmonary epithelial cells are the main cell type expressing ACE2, and are promptly infected by SARS-CoV-2. Furthermore, macrophages are strongly activated by the virus uptake^16^. Together, this leads to a strong immune activation upon infection, leading to the production of pro-inflammatory mediators^17,18^. Although the endothelial cell layer is likely only barely susceptible to SARS-CoV-2 infection, it is directly affected by the local inflammatory response^17^. This may occasionally lead to cell dysfunction, cell death and the deterioration of the barrier functions, which are characteristics of ARDS^17,19^.

Facing the prolongation of the COVID-19 pandemic and the appearance of new SARS-CoV-2 variants, waning immunity and vaccine hesitancy, an increase in the number of infections and admission in the intensive care units is being seen^1,20^. In severe cases of the disease, antiviral drugs may not be sufficient to ameliorate patient’s conditions, as a systemic inflammation may have already been established. The administration of glucocorticoids, targeting the uncontrolled hyper-inflammation has been evaluated on patients for its efficacy in reducing disease severity^21,22^. However, improvements were usually only reported for some specific cases^23,24^. Moreover, corticosteroids, like dexamethasone, were shown to induce lung epithelial cell senescence and promote pulmonary fibrosis, what might further contribute to lung tissue pathology^25,26^.

The production of cytokines and chemokines, which contribute to tissue inflammation and immune cell recruitment, is partially regulated by heat-shock proteins (HSPs)^27,28^. HSPs are one among the most abundant proteins in mammalian cells. They constitute a large family of chaperones that upon stress caused by heat, hypoxia or oxidative stress, assist the assembly, stabilization and translocation of proteins^27,29,30^. HSP90, including the isoforms HSP90α and HSP90β, is crucial for the maturation of proteins involved in cell division and differentiation, having a broad range of substrates, including kinases and transcription factors^27,31^. The pharmacological intervention through HSP90 inhibitors was shown to be beneficial in the most variated spectrum of diseases, including the treatment of cancer^32^, neurodegenerative diseases^33^, inflammatory diseases^28,34^ and infections^35^. The potent HSP90 inhibitor Ganetespib is being currently used in phase III studies in oncology and as an antiviral drug against Epstein-Barr virus (EBV) infection^36^. It has been shown to present promising pharmacokinetics properties^37^. HSP90 inhibition was also shown to be efficient in reducing viral replication *in vitro*, affecting the synthesis of important viral transcripts^35,38^. Moreover, they were shown to exert barrier protective functions on pulmonary arterial endothelial cells and possess important antiinflammatory properties. Therefore, being useful as a therapeutic strategy in ARDS and other pulmonary inflammatory diseases^34–36,39–41^. Wyler et al. (2020)^40^ showed an up-regulation of expression of hsp90 transcripts in Calu-3 and H1299 lung epithelial cells upon infection with SARS-CoV-2. Therefore, targeting HSP90 could be a valuable strategy to prevent disease progression and lung inflammation during severe cases of COVID-19.

The development of novel therapeutic strategies to better intervene in severe cases of COVID-19 is urgently needed. They should goal at controlling the hyper-inflammation generated locally and systemically and reducing viral replication. Here, we analyze the effects of the HSP90 inhibitor Ganetespib *in vitro* on infected alveolar epithelial cells (AECs) and alveolar macrophages, to characterize its effects on cell activation and viral replication. *In vivo*, in the Syrian hamster COVID-19 model, we evaluate its efficacy in controlling systemic inflammation and the viral burden after infection. *In vitro*, Ganetespib reduced viral replication on AECs in a dose-dependent manner and lowered significantly the expression of pro-inflammatory genes in both AECs and alveolar macrophages. *In vivo*, the administration of Ganetespib led to an overall improvement of the clinical condition of infected animals, with decreased systemic inflammation, reduced edema formation and tissue pathology.

## Material and Methods

### Ethic statement and COVID-19 hamster model

Experiments including female and male Syrian hamsters *(Mesocricetus auratus;* breed RjHan:AURA, JanvierLabs, France) were approved by the relevant state authority (LaGeSo) and executed in compliance with (inter-)national regulations. SARS-CoV-2 preparation and infection of hamsters were carried out as previously described^12^.

### Virus culture

SARS-CoV-2 isolate (BetaCoV/Germany/BavPat1/2020) was kindly provided by Daniela Niemeyer and Christian Drosten, from Charité Berlin, Germany. Virus stocks were propagated under BSL-3 conditions in Vero E6 cells (ATCC CRL-1586). Prior to infection, SARS-CoV-2 isolate was grown on Vero E6 cells in minimal essential medium (MEM; PAN Biotech) with 10% fetal bovine serum (PAN Biotech) and 100IU/ml Penicillin G and 100 μg/ml Streptomycin (Carl Roth). After incubation and titer determination, stocks were prepared and stored at −80°C. Integrity of the furin cleavage site was confirmed through sequencing of stocks prior to infection. *Cell culture*. Vero E6 cells (ATCC CRL-1586) were cultivated in Dulbecco’s modified Eagle’s medium (DMEM) supplemented with 10% heat-inactivated fetal calf serum (FCS), 1% non-essential amino acids, 1% L-glutamine and 1% sodium pyruvate (all Thermo Fisher Scientific), at 37 °C with 5% CO2. Alveolar epithelial cells were isolated from distal lung tissue and cultured as previously described^40^. Briefly, isolated primary cells were co-cultivated with gamma-irradiated mitotically inactivated NIH3T3 mouse embryonic fibroblasts (MEFs) in a 3:1 mixture of Ham’s F-12 nutrient mix (Life technologies) and DMEM supplemented with 5 % FCS, 0.4 μg/mL hydrocortisone (Sigma-Aldrich), 5 μg/mL recombinant human insulin (Sigma-Aldrich), 8.4 ng/mL cholera toxin (Sigma-Aldrich), 24 μg/mL Adenine (Sigma-Aldrich), 10 ng/mL recombinant human epidermal growth factor (Invitrogen), 0.1 μM Dibenzazepine (DBZ; Tocris) and 9 μM Y27632 (Miltenyi Biotec). Differentiation was induced by additional treatment with 3 μM CHIR-99021 (Sigma), 10 ng/ml keratinocyte growth factor (KGF; Invitrogen), 10 ng/ml fibroblast growth factor (FGF)-10 (Invitrogen), 100 μM 3-isobutyl-1-methylxanthine (Sigma), 100 μM 8-Bromoadenosine 3’,5’-cyclic monophosphate (Biolog), 25 nM Dexamethasone (Sigma) and 20 μM DBZ for 3 days. Two days prior to infection, the primary cells were separated from the MEFs by differential trypsinization and subsequently seeded in cell culture vessels in DMEM with 10% FCS, 1% non-essential amino acids, 1%, L-glutamine and 1% sodium pyruvate. Cells were washed with PBS and the same medium was added together with virus for 1 h at 37°C, with 5% CO_2_. Studies with human alveolar macrophages were approved by the local ethics committee. Alveolar macrophages were isolated from bronchoalveolar lavage performed for routine diagnostic purposes in patients, as previously described^42^. Cells were washed twice in cold PBS, then resuspended at 10^6^/ml in RPMI 1640/10 % FCS/penicillin/streptomycin. Untreated alveolar macrophages were placed into 24-well tissue culture plates and allowed to adhere for 2 h. Cell monolayer was washed three times to remove nonadherent cells and antibiotics and cultured in RPMI 1640/10 % FCS. Prior to infection, cell culture was washed three times with HBSS. Treatment with Ganetespib was done prior to infection with SARS-CoV-2 for characterization of its protective functions. AECs were treated with different concentrations of Ganetespib, ranging from 12.5 nM to 800 nM in a log2 scale and alveolar macrophages were treated for 16 h with 100 nM of this inhibitor.

### Animal model

To ensure successful intranasal infection, hamsters were anaesthetized with a triple anesthesia consisting of midazolam (2 mg/kg), butorphanol (2.5 mg/kg) and medetomidine (0.15 mg/kg). Subsequently 1×10^5^ PFU SARS-CoV-2 diluted in 60 μl MEM (PAN, Biotech) were applied intranasally in the hamsters^43^. In the first *in vivo* experimental set, Syrian hamsters were randomly assigned to 3 groups, all being infected. The intra-peritoneal (i.p.) treatment with 25 mg/kg body-weight of Ganetespib (kindly provided by Aldeyra Therapeutics) was performed together with infection at day 0 (group 1) or twice, both at day 0 and day 4 after infection (group 2). The third group received placebo therapy. In a second experimental set, animals were equally divided in 3 groups. The control group received placebo treatment at day 3 after infection, whilst the other two groups received Ganetespib at the time-point of 48 h or 72 h after infection. All animals included in the experiment were monitored for signs of disease twice-daily, with weight and temperature measured daily for compliance with score sheet criteria. Hamsters were selected randomly for all scheduled time-points for analysis. For euthanasia, animals were anesthetized with medetomidine (0.15 mg/kg), midazolam (2 mg/kg), and butorphanol (2.5 mg/kg) prior to cervical dislocation and exsanguination^43^. Serum, EDTA blood, oropharyngeal swabs and lungs were collected for virologic, histopathological and bulk sequencing analysis.

### Viral load quantification

Virus titers and RNA copies were determined by plaque assay and quantitative RT-PCR analysis as previously described^12^. RNA was extracted from oropharyngeal swabs and from 25 mg of homogenized lung tissue, with the innuPREP Virus DNA/RNA Kit (Analytic Jena, Jena, Germany), according to manufacturer’s instructions. For quantification of viral RNA, the NEB Luna Universal Probe One-Step RT-qPCR Kit (New England Biolabs) was used and the qPCR was conducted on the StepOnePlus Real-Time PCR System (Thermo Fisher Scientific)^44^. Titration was performed to quantify replication competent virus. Shortly, lung tissue was homogenized in a bead mill (Analytic Jena), and 50 mg of homogenized lung were used to prepare 10-fold dilutions starting from - 1 to - 6 and transferred onto Vero E6 cells seeded in 24-well plates (Sarstedt, Nümbrecht, Germany). Subsequently, the plates were incubated for 2,5 hours at 37°C and overlaid with 1,5% methylcellulose (Sigma Aldrich). After 72 h, cells were fixed in 4% formaldehyde and stained with 0.75% crystal violet. Results were evaluated by counting the plaques.

### Histopathology for SARS-CoV-2 infected hamsters

Lungs were embedded in paraffin and sections were made from tissue for immunostaining and analysis. Lung tissue pathology was evaluated by board-certified veterinary pathologists in a blinded fashion following standardized recommendations, including pneumonia-specific scoring parameters as described previously^12^.

### Bulk RNA analysis

Sequencing was performed after RNA isolation of the AECs and alveolar macrophages using Trizol reagent according to the manufacturer’s instructions. Libraries were constructed using the NEBNext Ultra II Directional RNA Library Prep Kit (New England Biolabs) and sequenced on a high throughput NextSeq 500 device. Reads were aligned to the M. auratus genome (MesAur1.0 downloaded from Emsembl and adapted as described previously^12^) using hisat2^45^. Gene expression was quantified using the package featureCounts from Rsubread^46^ and analyzed by DESeq2^47^. Differentially expressed genes were defined by an absolute fold change in mRNA abundance greater than 1.5 (log2 fold change of 0.58 - using DESeq2 shrunken log2 fold changes) and an adjusted p-value of less than 0.05 (Benjamini-Hochberg corrected).

### Mass spectrometry

Serum samples were mixed 1:1 with RIPA lysis buffer, while lung samples were prepared at a concentration of 25mg in 500 microliters of buffer. Samples were extracted by mixing 25 μL of sample with 25 μL of 25% w/w acetonitrile (ACN) in water, 100 μL of formic acid 0.1% and 400 μL of methyl-t-butyl ether (MTBE). Samples were vortexed and then centrifuged at 3000x g for 5 minutes. The supernatant was collected and samples extracted again with other 400 μL of MTBE. Both extracted supernatants were pooled dried under nitrogen and resuspended in 50 μL of 40% ACN in water before analysis. For the LC-MS/MS analysis, samples were analyzed on a Quantiva triple quadrupole (Thermo) coupled to a 1290 Infinity HPLC (Agilent), using a Zorbax Eclipse column (2.1 x 50 mm, 1.8 micron particle size). Separation was performed using a 0.3 ml/min gradient ranging from 40% of solvent B (acetonitrile with 0.1% formic acid; solvent A = water with 0.1% formic acid) to 95% in 2 minutes. The HESI source was operated with 3500 V spray voltage, 30 arbitrary units sheat gas, 8 arbitraty units auxiliary gas and 350 °C transfer tube and vaporizer temperature. Two transitions were monitored (365 -> 131 and 365 -> 323) in positive mode. Quantification was performed using QuanBrowser software (Thermo).

### Statistical analysis of hamster data

GraphPrism 9.1.2 software (GraphPad Software, USA) was used for statistical analysis of the clinical data and analysis of the transcripts of SARS-CoV-2 genome in the RNA sequencing. Details of the statistical analysis are described by each experiment at its respective figure legend. One-way or two-way ANOVA with Tukey’s multiple comparisons test for unpaired data was used for the *in vivo* data. Unpaired t-test was used to compare PFU counting on AEC supernatant. Further data on viral replication were compared using Mann-Whitney U test. P values are shown as following for significance: p<0.05 (*), p<0.01 (**).

## Results

### Ganetespib has antiviral and anti-inflammatory properties in AECs and alveolar macrophages

HSP90 inhibition was shown to hinder SARS-CoV-2 replication in Calu-3 lung epithelial cells and AECs^40,48,49^. To better characterize the antiviral and anti-inflammatory properties of Ganetespib on differentiated AECs, we infected cells for 1 h with 5 x 10^2^ PFUs of SARS-CoV-2, and treated them for 16 h with different concentrations of Ganetespib, ranging from 12.5 nM to 800 nM. Virus titer was assessed by the measurement of infectious viral particles in the AECs supernatant. Ganetespib treatment led to a statistically significant inhibition in viral production already at 100 nM. Effects on viral load reduction were similar in higher concentrations (Fig. 1A). Neufeldt and colleagues (2022)^50^ showed that upon infection of lung epithelial cells with SARS-CoV-2, NF-kB signaling was activated, leading to the expression of specific inflammatory cytokines. Contrarily, induction of interferon regulatory factors (IRFs) and interferon stimulated genes (ISGs) was not typically seen in COVID-19, probably due to the antagonist role of the open reading frame 6 (ORF6), an accessory gene of the SARS-CoV-2 genome, on interferon signaling^50–54^. To better characterize the impact of Ganetespib treatment on the expression of distinct inflammatory cytokines and stress-related genes, we performed RNA sequencing from infected AECs. Inhibition of HSP90 led to a substantial reduction of expression of NFKBIA transcripts. Moreover, transcripts of genes related to the IFN response, including IFIT2 and ISG15, and of several inflammatory cytokines, i.e. IFNB1 and CXCL10, typically up-regulated upon SARS-CoV-2 infection^11,19^, were overall reduced (Fig 1B). Interestingly, this reduction was already observed at a concentration of 12.5 nM Ganetespib and was more evident at higher concentrations, with expression values similar to that of uninfected cells. Expression of the stress-related genes ARRDC3 and DDIT3 was neither affected by SARS-CoV-2, nor by Ganetespib. Next, we assessed the expression of inflammatory cytokines and genes involved in the type I/II IFN antiviral response in infected AECs after treatment with 50 or 100 nM Ganetespib. Expression of pro-inflammatory cytokines and chemokines, including IL-6, IL-1b, CXCL10, CCL5 and TNFSF10, enhanced in COVID-19 patients^8,9,19^, was reduced after treatment with both Ganetespib concentrations. In addition, expression of NFKBIA, IFNy-related genes and genes induced by type I IFN response, including IFIT2 and IRF1, were reduced, when compared to control cells. Cellular stress markers, including JUN and FOS, also had reduced expression after treatment with Ganetespib (Fig. 1C).

**Figure 1:**
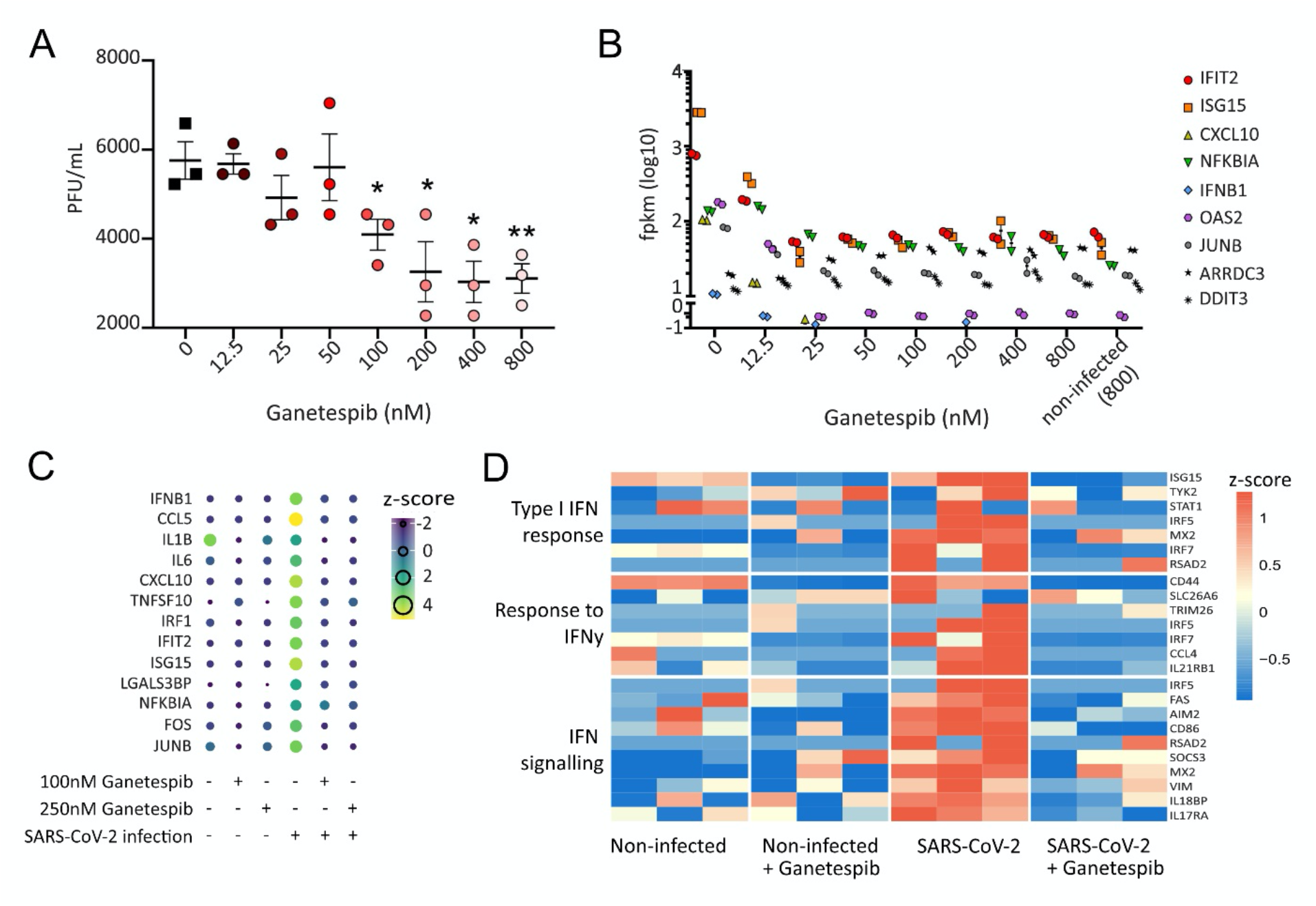
Antiviral and anti-inflammatory properties of Ganetespib in *in vitro* infected AECs and alveolar macrophages. AECs, isolated from human lung tissue, were infected with SARS-CoV-2 and treated with different concentrations of the HSP90 inhibitor Ganetespib. Concentrations ranged from 12.5 nM Ganetespib to a maximum of 800 nM on a log2 scale. Non-treated or non-infected cells serve as control. (A) Viral titers in supernatants of AECs infected with SARS-CoV-2 and treated with the indicated concentration of Ganetespib. (B) Expression analysis of selected induced genes in AECs after infection. Shown are normalized gene expression values compared to infected samples treated with DMSO of the same timepoints (normalized value after FPKM - fragments per kilo base of transcript per million mapped fragments). (C) Expression of selected genes in AECs after SARS-CoV-2 or mock-infection, and treatment with Ganetespib. Shown are DESeq2 normalized counts on a log10 scale. Alveolar macrophages were differentiated from lung tissue and infected with SARS-CoV-2 prior to treatment with Ganetespib. Mock infected cells, treated or non-treated, served as control. (D) Heatmaps of differently expressed genes in alveolar macrophages after infection with SARS-CoV-2 and treatment with Ganetespib. Uninfected cells, receiving Ganetespib or not, were used as controls. Genes related to IFN-mediated signaling pathway, type I IFN signaling and cellular-response to IFNy were selected, as previously described^57^. Columns represent samples and rows genes. Shown are z-scores of DESeq2-normalized data and color scale ranges from blue (10 % lower quantile) to red (10 % upper quantile) of the selected genes. N=3 in A, D, and n=2 in B and C. Unpaired t-test used for comparison in A. * p < 0.05, ** p < 0.01.

Monocytes and macrophages were suggested to be critical mediators of the inflammatory responses in COVID-19^9,55^. Inflammatory macrophages, and regulatory macrophages, which have been shown to have a pro-fibrotic nature, also play an important role in the hyper-inflammatory condition of COVID-19, contributing to tissue pathology^16,56^. To characterize the effect of Ganetespib on the expression of typical inflammatory cytokines and type I/II IFN antiviral response, infected alveolar macrophages were subjected to RNA sequencing. Similar to the observation made in infected AECs, genes belonging to the IFN response, including IRFs and ISGs, showed reduced expression after Ganetespib treatment. Ganetespib treatment reduced the expression of pro-inflammatory cytokines, including IL1b and IL6, genes belonging to IFN signaling and related to fibrosis, when compared to untreated cells (Fig. 1D and Supp. Fig. 1A).

### Administration of Ganetespib on infected hamsters does not affect pulmonary viral loads

Upon SARS-CoV-2 infection, Syrian hamsters present a moderate course of COVID-19, resembling many aspects of the disease in humans^12,43^. This model features high viral replication in the upper and lower respiratory tracts, strong immune cell influx in the lungs, pulmonary inflammation and lung tissue pathology^43^. To assess the effects of the HSP90 inhibitor Ganetespib as an antiviral and anti-inflammatory medicine in SARS-CoV-2 infection, Syrian hamsters were infected with 1 x 10^5^ PFU SARS-CoV-2 and divided into 3 groups with three animals each. Group 1 received Ganetespib at day 0, group 2 received Ganetespib at day 0 and at day 4 p.i. and a third untreated group served as control. Animals were sacrificed on days 3 (only the control group and group 1), 5, and 7 p.i. (Fig. 2A). Animals are depicted for their sex in each of the groups analyzed in Supp. Table 1. To monitor disease progression, body-weights of all animals were recorded daily. Infection with SARS-CoV-2 led to a progressive decrease of body-weight, as observed previously^12,43^. Administration of Ganetespib did not affect animal weight loss in any of the timepoints analyzed, with no significant differences between the treated and control groups (Fig. 2B). Viral loads were quantified from homogenized lung tissue (Fig. 2C) and oropharyngeal swabs (Fig. 2D) by RT-qPCR, and plaque assays were performed on the lung tissue homogenate (Supp. Fig. 2A). Virus was detected in both compartments and virus titers decreased over time. At day 7 p.i. replicating virus was undetectable in the lung tissue. Treatment with Ganetespib did not affect viral loads in lung tissue as determined by RT-qPCR and plaque assay. *In vitro*, we showed that the antiviral property of Ganetespib could be observed after treatment of the cells with concentrations higher than 100 nM. To quantify the systemic concentration of Ganetespib in the Syrian hamsters after treatment, mass spectrometry analysis was performed on lung homogenates and serum of the animals at days 1, 3, 5 and 7 after Ganetespib administration. At day 1 post-treatment, Ganetespib was detected in lungs with concentrations ranging from 10 to 15 nM, and in serum up to 6 nM. Ganetespib concentrations decreased progressively with time with almost non-detectable levels in both compartments at day 5 (Fig. 2E).

**Figure 2:**
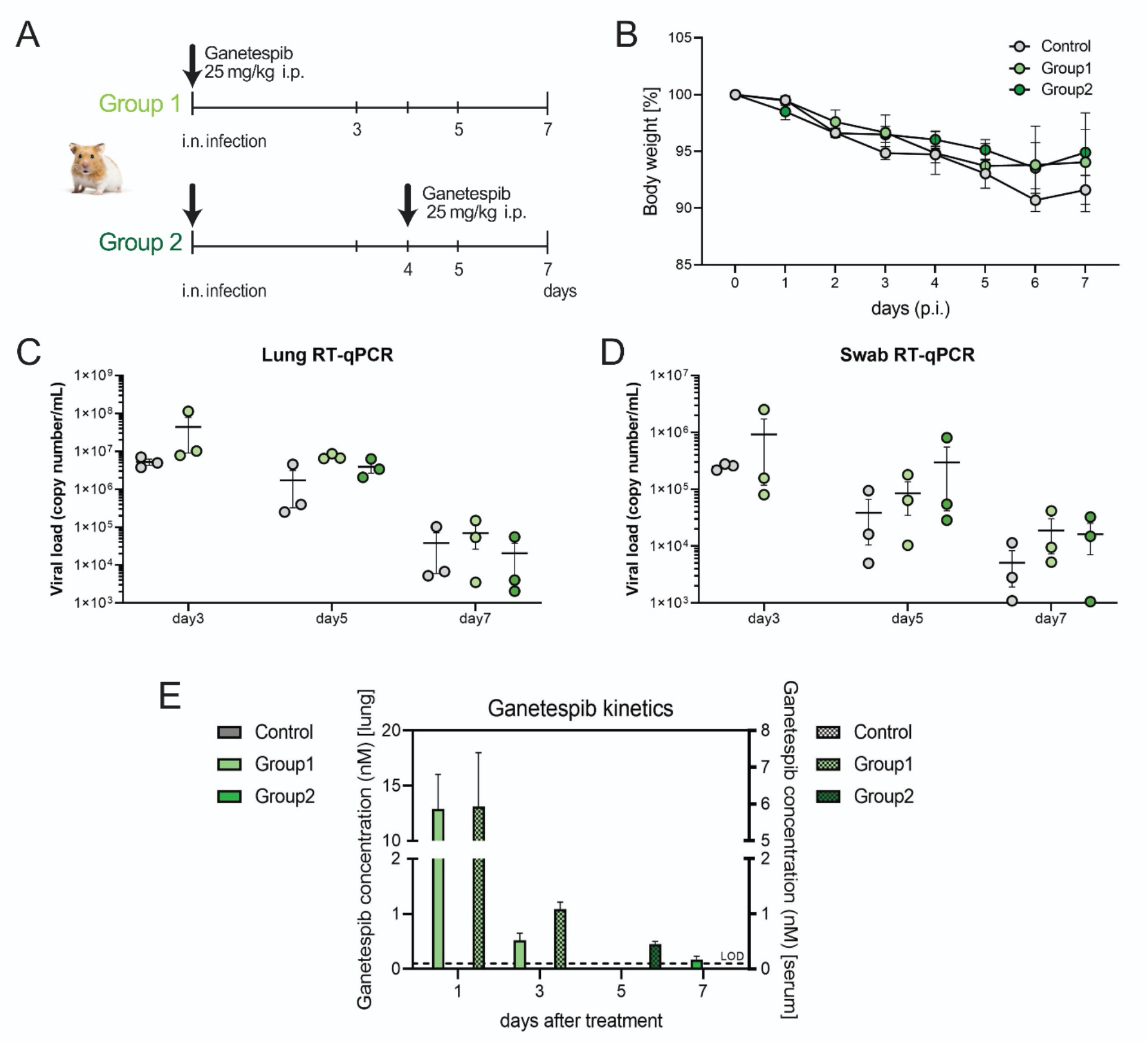
Treatment with Ganetespib does not affect viral loads in Syrian hamsters. (A) Syrian hamsters challenged with SARS-CoV-2 (1 x 10^5^ PFU), were treated with 25 mg/kg Ganetespib once, together with infection (group 1), or twice, at day 0 and at day 4 p.i. (group 2). Control group received same volume of dilutor at day 0 p.i. Analysis was made at day 3 (only for the control group and group 1), day 5 and day 7 p.i. (B) Body weight in percentage from the original weight was measured daily after infection of the animals with SARS-CoV-2. Viral load was quantified by RT-qPCR analysis of viral genomic RNA (gRNA) copies detected in homogenized lung tissue (C) and oropharyngeal swabs (D) at days 3, 5 and 7 p.i. (E) Pharmacokinetics of Ganetespib quantified by mass spectrometry in the lung tissue or serum of Syrian hamsters infected with SARS-CoV-2. Shown is the number of days after Ganetespib application (first or second doses). Results are displayed as mean ± SE and n=3. Comparisons were made with a one-Way ANOVA test.

### Ganetespib reduces lung tissue pathology in infected Syrian hamsters

Lung tissue histology was examined on longitudinal sections of the left lobes of the lungs of infected hamsters after treatment with Ganetespib and staining with hematoxylin and eosin. The lung pathology was assessed with a semi-quantitative scoring system to evaluate the protective role of Ganetespib after infection. Infected lung tissue presented increased infiltration of immune cells and accumulation of fluids with edema formation, seen by the darker areas on the histology of the whole tissue (Fig. 3A-F left). Analysis of the epithelial cell layer (Fig. 3A-F middle) and the alveolar space, including the endothelial cell layer (Fig. 3A-F right), showed increased cell hyperplasia and inflammation after infection. Treatment with Ganetespib was sufficient to reduce tissue pathology and inflammation, as seen in a general overview of the tissue, when compared to the untreated control group. Treatment with Ganetespib reduced pathology scores, particularly in the alveolar compartment with a significant reduction on the lung affected area among animals of group 2 at day 7 p.i. (Fig. 3G). The inflammation score was also reduced after treatment with Ganetespib, but not statistically different from the control group (Supp. Fig. 2B). The analysis of alveolar edema (Fig. 3H) and perivascular edema formation (Supp. Fig. 2C), presented an apparent, but not significant, reduction in the score levels after treatment with Ganetespib at day 5 and 7 p.i., suggesting a reduction in endothelial layer damage and lung barrier dysfunction. Reduced alveolar epithelial type 2 (AT2) cell hyperplasia and endotheliitis (Fig. 3I and J) was observed after treatment at day 5 p.i., in comparison to the control group, but only a reduction in endotheliitis among animals receiving twice Ganetespib was statistically different from the control group. The pathology score of alveolar epithelial necrosis (Supp. Fig. 2D), bronchitis (Supp. Fig. 2E) and immune cell infiltration, including lymphocytes, macrophages and neutrophils (Supp. Fig. 2G-I), showed no differences among treated and untreated animals. However, Ganetespib treatment led to a significant decrease in broncho-epithelial hyperplasia among animals treated twice with Ganetespib, compared to the control group (Supp. Fig. 2F).

**Figure 3:**
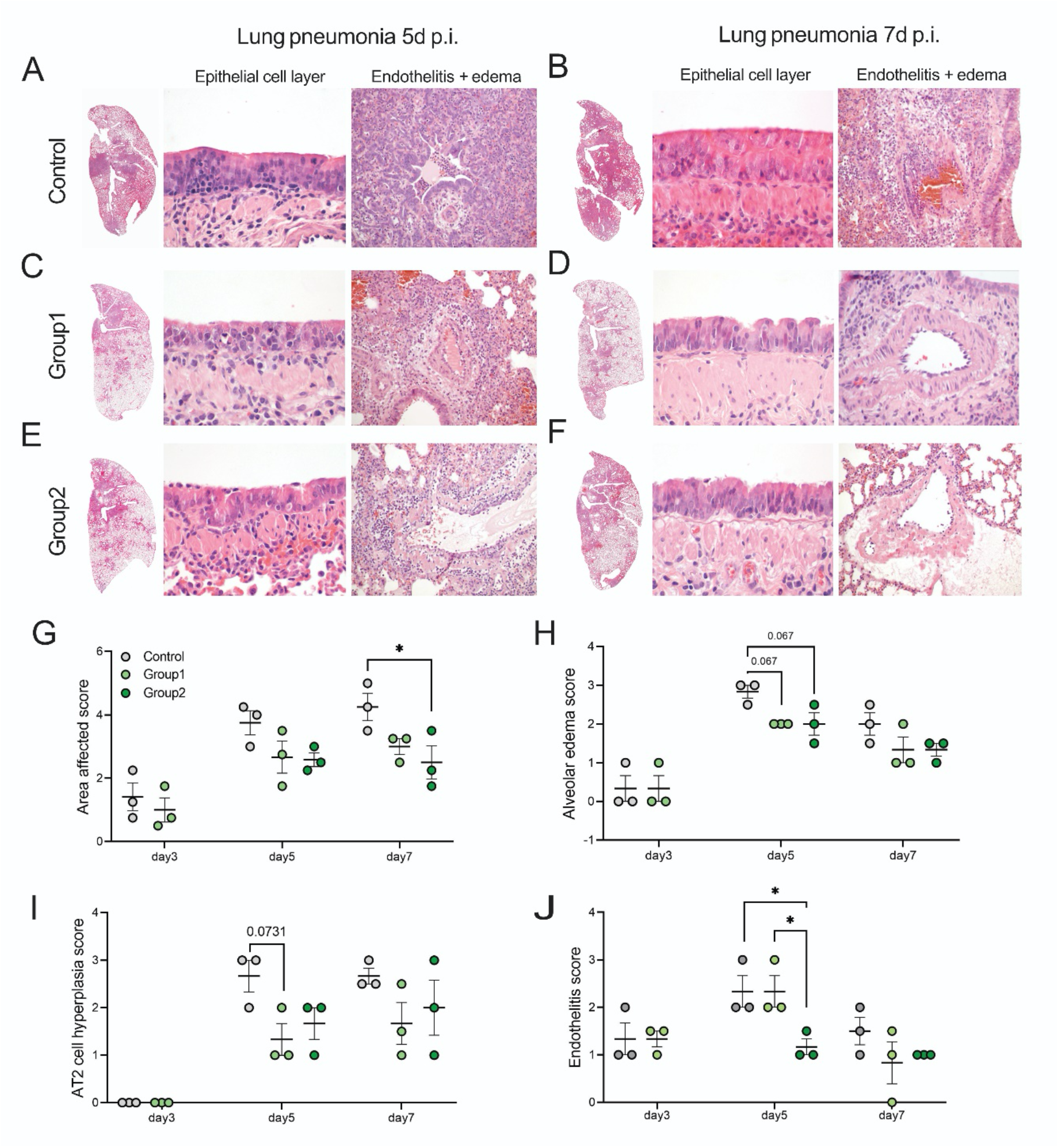
Lung tissue pathology scores are reduced in infected hamsters after treatment with Ganetespib. Longitudinal sections of left lungs from Syrian hamsters after 5- or 7-days infection with SARS-CoV-2 and receiving treatment of Ganetespib. Treatment was performed at day 0 p.i. or at day 0 and again at day 4 p.i. and lung tissue was stained with hematoxylin and eosin for histological comparisons. Shown are whole lung sections, and magnification of the epithelial or endothelial layers. Lung pneumonia with edema formation is seen in darker colors, due to the accumulation of fluids and cells. Representative histology is shown. Semi-quantitative analysis of histological lesions quantified at day 3, 5 and 7 p.i. were scored and shown for comparisons between the groups. Lung affected area (G), alveolar edema (H), AT2 cell hyperplasia (I) and lung endotheliitis (J) are shown. Results are displayed as mean ± SE and n=3. Comparisons were made with a one-Way ANOVA test. * P < 0.05.

### Ganetespib reduces lung inflammation in infected Syrian hamsters also when applied at later time-points

In our animal model for COVID-19, the disease course is relatively fast when compared to human patients, with animals presenting pronounced disease symptoms already 2-3 days p.i. In order to better translate the actual treatment course in human patients, we repeated the infection model of Syrian hamsters with a later application time-point of the HSP90 inhibitor. Infected hamsters were divided into 3 groups with four animals each, receiving mock treatment (GB control), or treatment with Ganetespib at day 2 (GB 48h) or at day 3 p.i. (GB 72h), and animals were analyzed on day 5 p.i. (Fig. 4A). Sex of the animals analyzed is depicted in supp. table 1. Animal body-weight changes were recorded daily after infection and already at day 2 p.i. a decrease in body-weight was seen in all groups, reaching a maximum of 5 % reduction to the original weight at day 5 p.i.. No significant weight loss differences were observed after treatment of the infected animals with Ganetespib, compared to the control group (Fig. 4B). The concentration of Ganetespib after treatment in lung homogenate and serum was measured by mass spectrometry at day 5 p.i., which represented day 3 (GB 48h) or 2 (GB 72h) after Ganetespib application. Ganetespib concentrations in the lung were below 10 nM 2 days after Ganetespib application (GB 72h), similar to what was seen before, and slightly lower 3 days after application.

**Figure 4:**
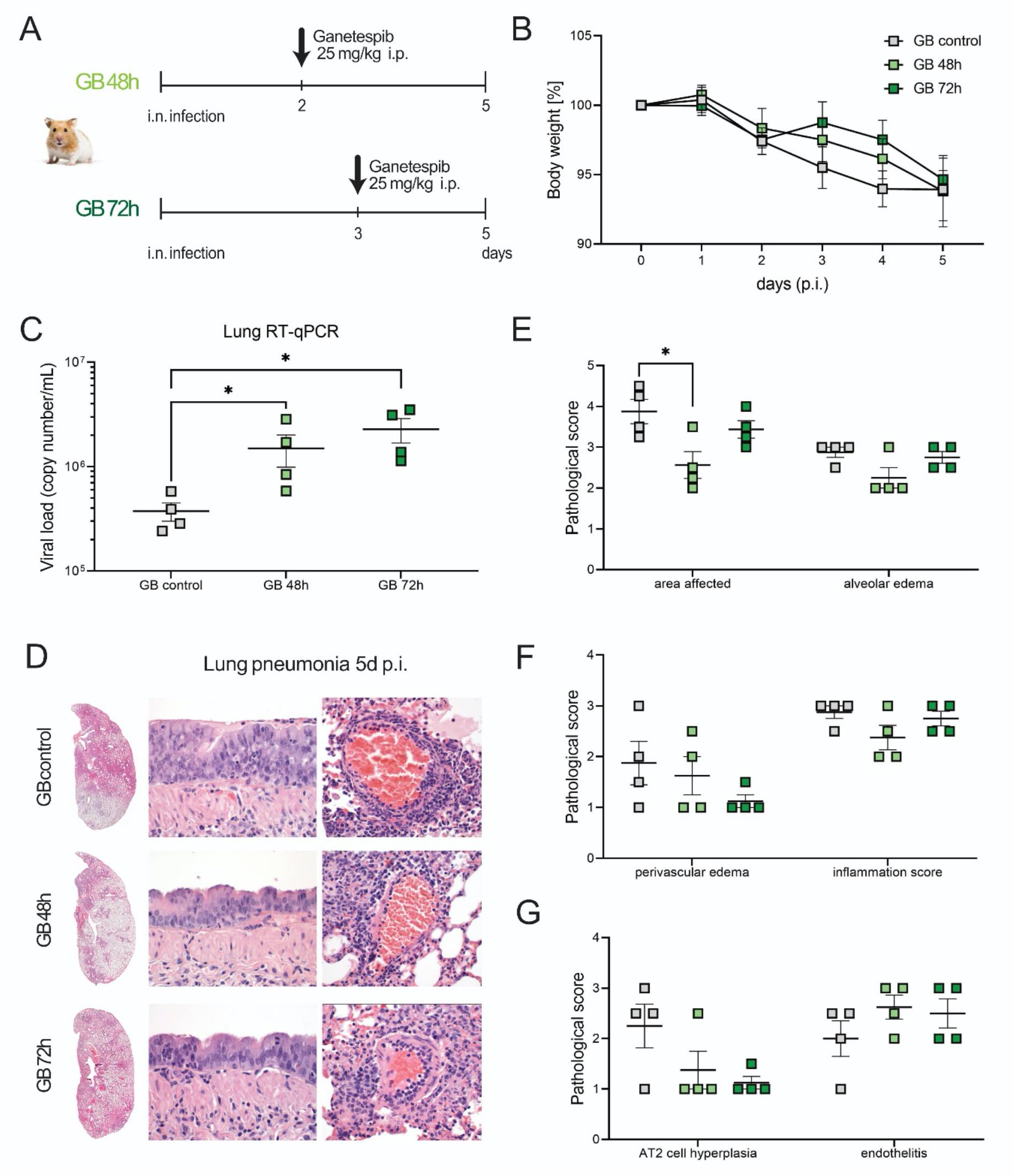
Early treatment with Ganetespib reduces inflammation and lung tissue pathology. (A) Syrian hamsters were challenged with SARS-CoV-2 (1 x 10^5^ PFU) and treated with 25 mg/kg Ganetespib at 48 h p.i. (GB 48h) or at 72 h p.i. (GB 72h). Control group received same volume of dilutor and analysis was made at day 5 p.i. (B) Body weight in percentage from the original weight was measured daily after infection of the animals with SARS-CoV-2. (C) RT-qPCR analysis of viral genomic RNA (gRNA) copies detected in homogenized lung tissue was performed at day 5 p.i. (D) Longitudinal sections of left lungs of Syrian hamsters 5 days after infection with SARS-CoV-2 were stained with hematoxylin and eosin for histological comparisons. Shown are whole lung sections, and sections focusing on epithelial cell layer and interstitial space. Lung pneumonia with edema formation is seen in darker colors, due to accumulation of fluids and cells. Representative histology is shown. Semi-quantitative analysis of histological lesions quantified at day 5 p.i. were scored and shown for comparisons between the groups for lung affected area and alveolar edema (E), perivascular edema and inflammation score (F) and AT2 cell hyperplasia and endotheliitis (G). Shown is the comparison between the groups receiving Ganetespib at day 2 or 3 p.i. and the control group. Average values ± SE are shown and n=4. Comparisons were made with a two-tailed Mann-Whitney test (C) and a one-Way ANOVA test (E-G). * P < 0.05.

In serum, similar values were observed among animals receiving Ganetespib 48h p.i., while in the second group it was not detectable (Supp. Fig. 3A). RT-qPCR analysis of the lung tissue for viral genome shows an increase in the viral burden in both animal groups treated with Ganetespib, in comparison to mock-treated animals (Fig. 4C). Viral titers of the group GB 72h were slightly higher than the titers of the GB 48h group, but not statistically different. This increase in viral titers was not observed by analysis of the lungs by plaque assay (Supp. Fig. 3B) or by RT-qPCR of swabs of infected animals (Supp. Fig. 3C). Histology of the lung tissue was also performed after staining with hematoxylin and eosin for characterization of tissue damage and cellular infiltration (Fig. 4D). Overall, the area of the lungs affected by the infection was reduced with the treatment at 48 h p.i., compared to control, with decreased edema formation, epithelial and endothelial cell damage, not seen among the animals treated with Ganetespib at 72h p.i.. Analysis by semi-quantitative scores of lung tissue pathology showed a significant reduction in the total area affected by pneumonia after treatment with Ganetespib at 48 h p.i. when compared to the control group. Alveolar edema formation was also reduced among the animals receiving Ganetespib at 48 h p.i., but not significantly different to the control group (Fig. 4E). The perivascular edema formation and inflammation scores on the lungs (Fig. 4F), AT2 cell hyperplasia and endotheliitis (Fig. 4G), alveolar epithelial necrosis, bronchitis and broncho-epithelial hyperplasia (Supp. Fig. 3D) and immune cell infiltration (Supp. Fig. 3E) were not affected by the treatment with Ganetespib, when compared to the control group, with only an overall tendency in reduction seen after treatment.

## Discussion

Facing the prolongation of the COVID-19 pandemic and the unequal level of vaccination around the globe, the appearance of new SARS-CoV-2 variants due to adaptations of the virus to evade humoral immunity is being seen^1,20^. Aiming at protecting the more vulnerable patients that may develop severe cases of the disease, different treatment strategies that aim at reducing virus titers and the systemic inflammation taking place after infection are urgently required. Here, we demonstrate the positive effects of Ganetespib, an HSP90 inhibitor, in reducing SARS-CoV-2 titers on AEC cells *in vitro* and its anti-inflammatory and tissue protective role in a hamster model of SARS-CoV-2 infection. *In vitro*, Ganetespib could reduce viral loads in AECs and led to a reduced expression of important pro-inflammatory cytokines on AECs and alveolar macrophages, which are normally associated with COVID-19. *In vivo*, we showed that, though Ganetespib had limited antiviral capacities, lung tissue pathology severity was reduced by the treatment with this inhibitor. Inflammation area, edema formation and lung epithelial cell damage were overall reduced after treatment of infected hamsters with Ganetespib, evidencing the positive properties of Ganetespib as a potential medicine for moderate and severe cases of COVID-19.

Many viruses were shown to be highly dependent on HSP90 for replication, depending on this chaperone for proper folding and function of the newly rapidly-synthesized viral proteins^38,58^. HSP90 inhibition by Ganetespib regulates viral replication by affecting the degradation or deactivation of important proteins involved in the replication machinery, such as particle assembly, exosome formation and cellular infection^35,59^. Therefore, inhibition of HSP90 can lead to a reduction in viral replication, as shown for HSV type I and EBV^35,39,60^. Here, we show that upon infection of AECs by SARS-CoV-2, Ganetespib was efficient in reducing viral loads in these cells in a dose-dependent manner, with effects observed with concentrations above 100 nM. Contrarily to AECs, we did not observe a reduction in viral loads in infected hamsters after treatment with Ganetespib, where rather an increase in the viral load was seen. Different from the *in vitro* experiment, the concentration of the HSP90 inhibitor Ganetespib achieved *in vivo* after i.p. application was considerably low, being comparable to the lowest doses analyzed *in vitro*. In our hamster model, concentrations achieved in serum and lung tissue were approximately 10 nM after 1-day application, with reduced or non-detectable levels at later time-points, as seen by the mass spectrometry analysis. Normal ranges of *in vitro* applications vary between 100 nM and 1 μM^36,40^. However, even if not sufficient to block viral replication, the systemic levels of Ganetespib showed efficient anti-inflammatory properties, dampening the production of important cytokines and chemokines after infection, as described by others^12,52,61^. With increased viral titers and reduced inflammation, an impaired immune response to the virus was seen. Future experiments aiming at the HSP90 inhibition should better consider the pharmacokinetics of this molecule, and consider distinct mechanisms to provide higher doses of this inhibitor systemically, possibly through distinct routes of application.

The pleiotropic effects of HSP90 include degradation of distinct proteins, including MAPK, JAK/STAT and NF-kB, affecting distinct signaling pathways implicated in inflammation, cell proliferation and fibrogenesis^27,62–64^. HSP90 inhibition was shown to promote the reduction of oxidative stress and inflammation of the lung tissue, and in this way, influence the recruitment and activation of important immune cells, including neutrophils and macrophages^28^. Production of important pro-inflammatory cytokines, such as IL-1β, IL-6, CXCL10, TNFSF10 and TNF, which are correlated to the progression of COVID-19, were shown to be reduced by HSP90 inhibition, probably due to the role of HSP90 in the IKK and NF-kB activation pathway^40^. Here, we could also show that Ganetespib could abrogate NF-kB signaling, with reduction in the expression of NFKBIA and cytokines after treatment. Impaired type I IFN immunity is a major risk factor in COVID-19, and is normally associated with the inadequate immune response observed in aged individuals. IFN signaling was shown to have a pivotal role in COVID-19 by coordinating an appropriate inflammatory response against viral infection and the subsequent activation of the innate and adaptive immune response ^65,66^. We could also observe a reduction in the expression of important genes involved in the IFN-signaling pathway, that may have contributed to the reduced tissue pathology observed. It is possible that the overall reduction of pro-inflammatory mediators observed could have led to a decreased activation of immune cells, affecting the animal immune response to the infection and further contributing to the higher viral titers observed.

Previous studies by our group showed a strong effect of the glucocorticoid Dexamethasone, applied alone or in combination with a monoclonal antibody targeting SARS-CoV-2, in modulating the immune response against virus infection^52^. Although this glucocorticoid was promising in reducing disease severity in both Syrian and Roborovski dwarf hamsters (this model approaching the severe cases of the disease in human patients), it showed significant effects in reducing the immune cell infiltration into the infected tissue. This could lead to an exacerbated viral spread in the tissue and the prolonged activation of the immune system, contributing to the development of different pathological conditions in the patient, including post-COVID-19 lung fibrosis. In contrast to Dexamethasone administration, Ganetespib had no adverse effect on the immune cell compartment, necessary for the viral control on the lung tissue, as we could not detect any differences in cell counts observed in the lung tissue after treatment.

Reports have characterized the development of fibrotic lungs in COVID-19 patients, which may progress even after disease resolution, with virus eradication^67–69^. Monocytes and profibrotic macrophages play a critical role in the pathogenesis and progression of both ARDS and lung fibrosis^70–72^ and a pronounced infiltration of monocyte-derived macrophages, which have a profibrotic nature, was seen in the lung of severe COVID-19 patients^16^. Lung fibrosis is driven predominantly by TGF-β, which normally is involved in tissue repair and resolution of the damage, and an exacerbated TGF-β production is frequently observed in COVID-19^10,73^. Inhibition of HSP90 was shown to be effective in attenuating fibroblast activation and blocking the signaling pathway of TGF-β, inhibiting the production of extracellular matrix and collagen^27,74,75^. Ganetespib could likely attenuate the progression of fibrosis in a mouse model of pulmonary fibrosis when applied daily for 3 weeks, suggesting a role in the treatment of fibrotic lung disease, especially problematic in ARDS patients receiving mechanical ventilation^74^. Here, we fail to show a reduction of fibrotic markers after treatment of alveolar macrophages with Ganetespib. Further experiments are needed to better characterize the development of fibrotic lungs in COVID-19 and to evaluate the role of Ganetespib as possible anti-fibrotic therapeutic.

The endothelial cell layer, in contrast to epithelial cells, has low expression of the ACE2 receptor. Infection of these cells by SARS-CoV-2 occurs therefore mainly via uptake of viral particle by endocytosis, not contributing to viral replication. However, upon tissue infection, endothelial cells can also get activated and contribute to the inflammatory responses^17^. The anti-inflammatory property observed by HSP90 inhibition was shown to protect the endothelial layer, contributing to a reduction in barrier dysfunction^76^. Here, we also show that Ganetespib has a protective role on the endothelial cell layer, as seen by reduced lung tissue endotheliitis, which may have contributed to a decrease in lung barrier dysfunction and edema formation. However, a delayed application of Ganetespib (at 72h p.i.) was not sufficient to protect animals from disease progression, with lung tissue pathology being similar to the observed among untreated animals. Probably, a delayed treatment with Ganetespib, combined with an increase in viral loads had the opposite effect to what was expected, with increased and prolonged inflammation of the tissue. To better determine the efficacy of this inhibitor in reducing tissue inflammation and tissue pathology, frequency and timing of the administration of this inhibitor should be taken into consideration.

Altogether, we describe here distinct beneficial properties of HSP90 inhibition with Ganetespib in controlling cellular activation and tissue inflammation, and in this line, contributing to tissue protection in COVID-19. We postulate that in future experiments, a combinatory strategy of treatment with Ganetespib and different antiviral compounds could be beneficial in moderate and severe cases of the disease, where Ganetespib would promote stabilization of the endothelial barrier of the lung, diminish inflammation and promote tissue protection, whilst reduced viral replication would lead to improvements in the disease outcome.

## Acknowledgements

We thank the support of Jeannine Wilde and Madlen Sohn from the BIH/MDC Genomics Technology platform in Berlin for the sequencing. LGTA, EW and ML are supported by the Project “Virological and immunological determinants of COVID-19 pathogenesis – lessons to get prepared for future pandemics (KA1-Co-02 ‘COVIPA’)”, a grant from the Helmholtz Association Initiative and Networking Fund. ACH was supported by BMBF (RAPID). KH and ACH were funded by BMBF (NUM-COVID 19, Organo-Strat 01KX2021), Charite 3^R^, Einstein Foundation EC3R as well as by DFG (SFB-TR84).

## Supplementary Data

**Supp. Figure 1:**
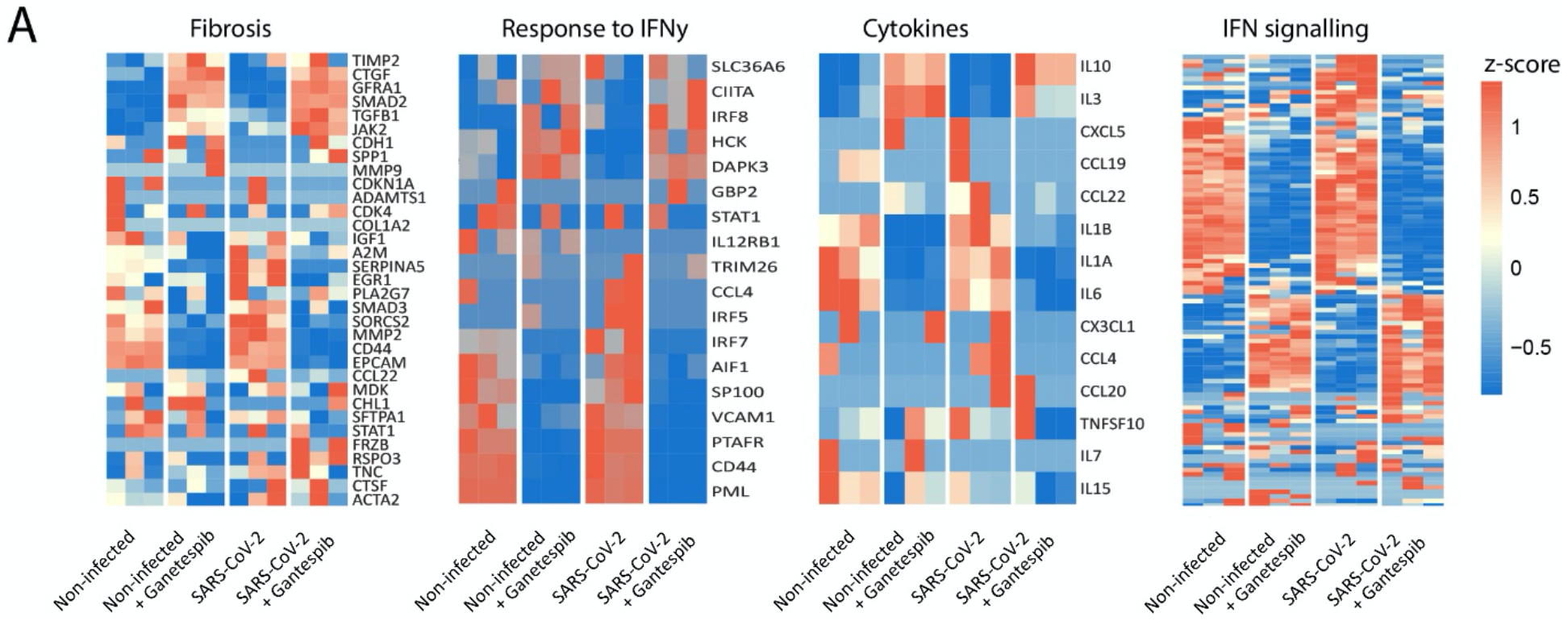
Anti-inflammatory properties of Ganetespib. Alveolar macrophages were isolated from human lung tissue and cultured in medium prior to the treatment with 100 nM of the HSP90 inhibitor Ganetespib. Non-treated or non-infected cells serve as control. (A) Heatmaps of differently expressed genes in alveolar macrophages after infection with SARS-CoV-2 and treatment with Ganetespib. Genes related to fibrosis response, the cellular-response to IFNy, cytokines and the IFN-mediated signaling pathway were selected, as previously described^57^. Columns represent samples and rows genes. Shown are z-scores of DESeq2-normalized data and color scale ranges from blue (10 % lower quantile) to red (10 % upper quantile) of the selected genes. N=3.

**Supp. Table 1:** Animal cohort information. Sex of animals used for both *in vivo* experiments in the respective groups analyzed.

|  | Experiment 1 |  |  |  |  |  |  |  |  | Experiment 2 |  |  |
| --- | --- | --- | --- | --- | --- | --- | --- | --- | --- | --- | --- | --- |
|  | Control |  |  | Group1 |  |  | Group2 |  |  | Control | GB 48h | GB 72h |
| Days p.i. | 3 | 5 | 7 | 3 | 5 | 7 | 3 | 5 | 7 | 5 | 5 | 5 |
| Male | 2 | 1 | 2 | 1 | 2 | 1 | - | 1 | 2 | 2 | 2 | 2 |
| Female | 1 | 2 | 1 | 2 | 1 | 2 | - | 2 | 1 | 2 | 2 | 2 |

**Supp. Figure 2:**
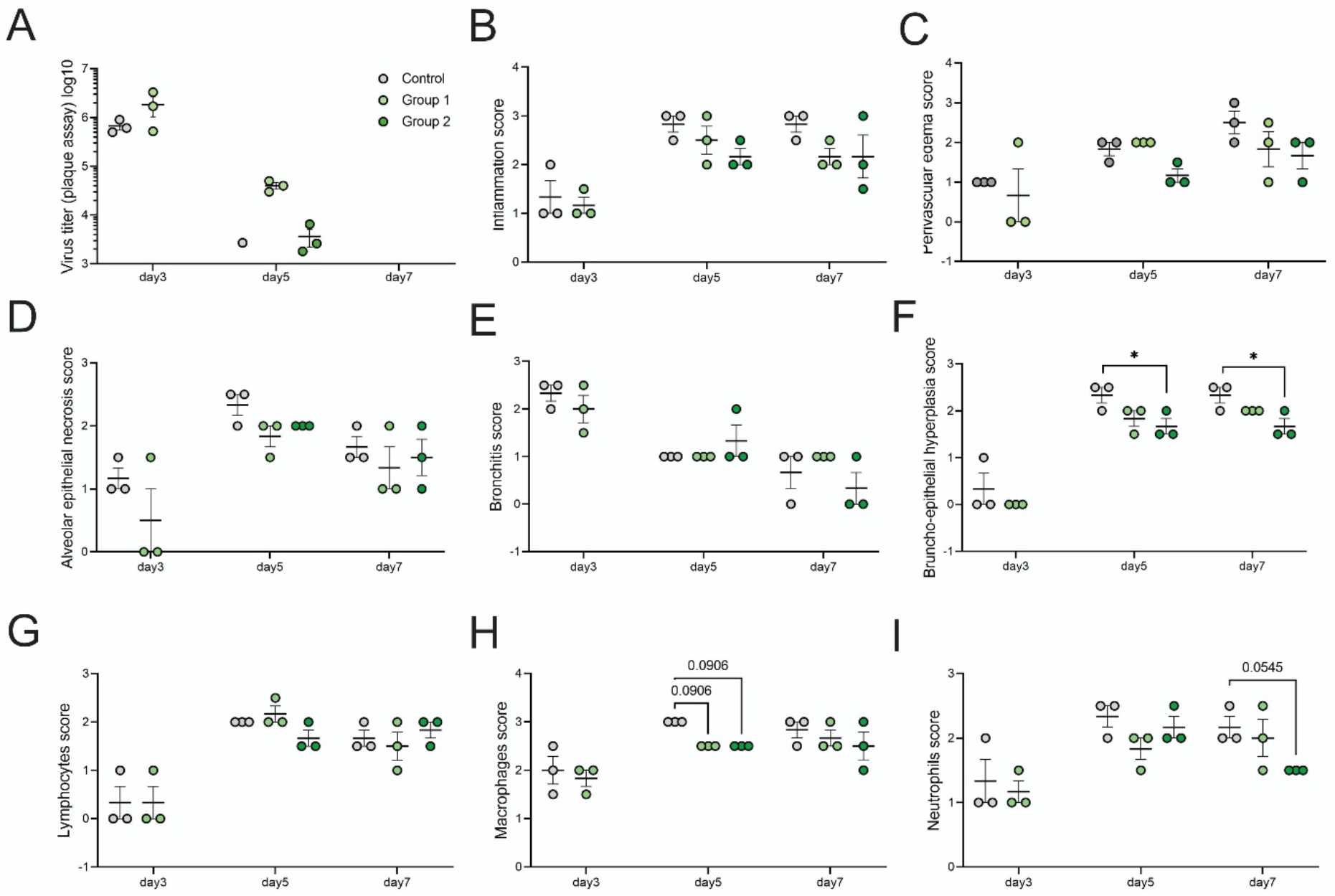
Lung tissue pathology scoring is improved after early treatment with Ganetespib. (A) Quantification of replication competent virus as plaque-forming units (PFU) per gram homogenized lung tissue. Longitudinal sections of left lungs from Syrian hamsters infected with SARS-CoV-2 and receiving a prophylactic treatment with Ganetespib at day 0 or day 0 + 4 p.i. after 3-, 5- or 7-days infection were evaluated by a semi-quantitative pathological scoring. Shown are scores for inflammation (B), perivascular edema (C), alveolar epithelial necrosis (D), bronchitis, (E), broncho-epithelial cell hyperplasia (F), and infiltration of lymphocytes (G), macrophages (H) and neutrophils (I). Results are displayed as mean ± SE and n=3. Comparisons were made with a two-Way ANOVA test. * P < 0.05.

**Supp. Figure 3:**
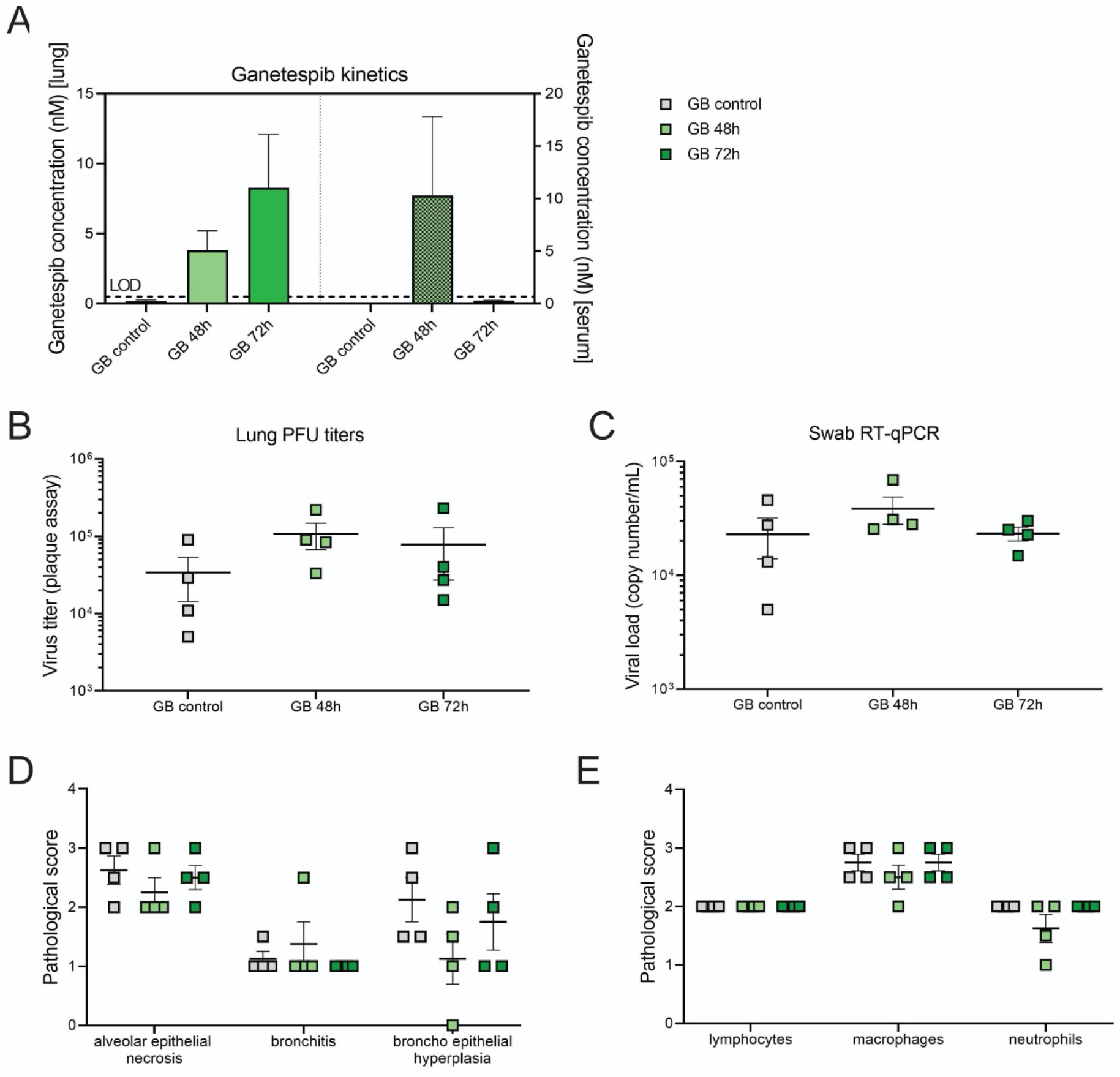
Treatment with Ganetespib at 48h p.i. reduces inflammation in the lungs in SARS-CoV-2 infected-hamsters. Syrian hamsters were challenged with SARS-CoV-2 (1 x 10^5^ PFU) and treated with 25 mg/kg Ganetespib at 48 h p.i. (GB 48h) or at 72 h p.i. (GB 72h). Control group received same volume of dilutor and analysis was made at day 5 p.i. (A) Pharmacokinetics of Ganetespib quantified by mass spectrometry in the lung tissue or serum of Syrian hamsters after 3 (GB 48h) or 2 days (GB 72h). Concentrations are shown in nM. (B) Quantification of replication competent virus as plaque-forming units per gram homogenized lung tissue was analyzed for viral titers. (C) RT-qPCR analysis of viral genomic RNA (gRNA) copies detected in oropharyngeal swabs at day 5 p.i. Semi-quantitative analysis of histological sections of the left lungs from hamsters analyzed after 5-days infection were scored and compared for the pathological score of alveolar epithelial necrosis, bronchitis and broncho-epithelial cell hyperplasia (D), and for immune cell infiltration (E), as shown. Results are displayed as mean ± SE and n=4.

